# Dementia risk factors modify hubs but leave other connectivity measures unchanged in asymptomatic individuals: a graph theoretical analysis

**DOI:** 10.1101/2020.10.08.331025

**Authors:** Hannah Clarke, Eirini Messaritaki, Stavros I Dimitriadis, Claudia Metzler-Baddeley

**Author notes:** **<u>Corresponding Author</u>** Claudia Metzler-Baddeley, Cardiff University Brain Research Imaging Centre (CUBRIC), Maindy Road, Cathays, Cardiff, CF24 4HQ, UK. /.

## Abstract

**Background:** Alzheimer’s Disease (AD) is the most common form of dementia with genetic and environmental risk contributing to its development. Graph theoretical analyses of brain networks constructed from structural and functional MRI measurements have identified connectivity changes in AD and individuals with mild cognitive impairment (MCI). However, brain connectivity in asymptomatic individuals at risk of AD remains poorly understood.

**Methods:** We analysed diffusion-weighted magnetic resonance imaging (dMRI) data from 160 asymptomatic individuals (38-71 years) from the Cardiff Ageing and Risk of Dementia Study (CARDS). We calculated white matter tracts and constructed whole-brain, default-mode-network and visual structural brain networks that incorporate multiple structural metrics as edge weights. We then calculated the relationship of three AD risk factors, namely Apolipoprotein-E ε4 genotype (APOE4), family history (FH) of dementia, and central obesity, on graph theoretical measures and hubs.

**Results:** We observed no risk-related differences in clustering coefficients, characteristic path lengths, eccentricity, diameter and radius across the whole-brain, default-mode-network or visual system. However, a hub in the right paracentral lobule was present in all high-risk groups (FH, APOE4, obese) but absent in low-risk groups (no FH, APOE4-ve, healthy weight).

**Discussion:** We identified no risk-related effects on graph theoretical metrics in the structural brain networks of cognitively healthy individuals. However, high-risk was associated with a hub in the right paracentral lobule, an area with motor and sensory functions related to the lower limb. If this phenotype is shown to predict symptom development in longitudinal studies, it could be used as an early biomarker of AD.

**Impact Statement:** Alzheimer’s Disease is a common form of dementia which to date has no cure. Identifying early biomarkers will aid the discovery and development of treatments that may slow AD progression in the future. In this paper we report that asymptomatic individuals at heightened risk of dementia due to their family history, Apolipoprotein-E ε4 genotype and body adiposity have a hub in the right paracentral lobule which is absent in low-risk groups. If this phenotype were to predict the development of symptoms in a longitudinal study of the same cohort, it could provide an early biomarker of disease progression.

## Introduction

Alzheimer’s disease (AD) is one of the major causes of dementia that affects 10% of individuals over the age of 65. In the United States, over one million individuals per year will be affected by AD by 2050 (Hebert et al., 2013). A recent review by the Lancet Commission concluded that almost half of dementia cases might be prevented or delayed by modifying 12 risk factors (Livingston et al., 2020). It emphasized the importance of improving the early detection of individuals at risk of developing AD so that preventative therapeutics can be discovered and developed in the future. It is therefore important to gain a better understanding of how AD risk factors affect the structure of the brain in healthy individuals and how risk-related effects differ from those of healthy aging.

The human brain has been characterized as a network of cortical and subcortical areas (network nodes) which communicate with each other via white matter tracts (connections, or edges) that carry neuronal signals (E. T. Bullmore & Bassett, 2011; Rubinov & Sporns, 2010). Structural networks can be derived from diffusion-weighted magnetic resonance imaging (dMRI) data via tractography (Basser et al., 2000; Mukherjee, Berman, et al., 2008; Mukherjee, Chung, et al., 2008), and are represented mathematically by graphs. Graph theory can then be employed to quantify the local and global organizational properties of the brain’s structural connectome (E. Bullmore & Sporns, 2009). Graph theoretical analyses of brain networks have provided insight into the effect of AD on the brain’s connectivity (Dai et al., 2019; John et al., 2017; Lo et al., 2010). More specifically, there is strong evidence that, even though AD pathology is initially present in localized brain areas, it still affects the whole brain as a network. It is, therefore, possible that people at risk of developing AD could show alterations in their structural brain networks and their graph theoretical metrics before developing the disease. This implies that investigations into possible relationships between AD risk factors and graph theoretical metrics of structural brain networks could provide biomarkers that signal disease onset or track disease progression.

In the present study, we used graph theory to characterize the mesoscale of structural brain networks for the whole-brain connectome and for systems that are affected in AD, namely the default mode network (DMN), as well as the visual network as a control (Badhwar et al., 2017), in 165 cognitively healthy individuals from the Cardiff Ageing and Risk of Dementia Study (CARDS) (38-71 years) (Coad et al., 2020; Metzler-Baddeley, Mole, Leonaviciute, et al., 2019; Metzler-Baddeley, Mole, Sims, et al., 2019; Mole, Fasano, Evans, Sims, Hamilton, et al., 2020; Mole, Fasano, Evans, Sims, Kidd, et al., 2020) with different risk factors for AD. The risk factors investigated were Apolipoprotein-E ε4 (APOE4), family history of dementia (FH) and central obesity. A statistical framework was followed to reveal potential differences in structural network organization between groups of aggregated risk levels. Our hypothesis was that individuals at the highest risk of dementia, i.e. obese APOE4 carriers with a family history of dementia, compared to those at lowest risk, i.e. normal-weighted non-carriers without a family history, would have altered integration and segregation parameters (increased characteristic path lengths, decreased clustering etc). In our exploratory analysis of hubs, we aimed to identify any highly interconnected nodes which consistently changed – gained or lost – in the transition from a low to high-risk group (FH vs. no FH, APOE4 carrier vs. non-carrier, obese vs. healthy weight).

## 1. Materials and Methods

Details of the CARDS study have been previously published (Coad et al., 2020; Metzler-Baddeley, Mole, Leonaviciute, et al., 2019; Metzler-Baddeley, Mole, Sims, et al., 2019; Mole, Fasano, Evans, Sims, Hamilton, et al., 2020; Mole, Fasano, Evans, Sims, Kidd, et al., 2020) and will hence only be briefly described in the following. The CARDS study was approved by the School of Psychology Research Ethics Committee at Cardiff University (EC.14.09.09.3843R2) and all participants provided written informed consent.

### 1.1 Participants

Individuals between the ages of 38 and 71 were recruited from the local community via Cardiff University community panels, notice boards and poster advertisements. Exclusion criteria included a history of neurological and/or psychiatric disease, severe head injury, drug or alcohol dependency, high risk cardio-embolic source or known significant large-vessel disease. MRI screening criteria were fulfilled by 166 participants. Table 1 summarises their demographic background, and information about their genetic and lifestyle risk variables. Depression was screened for with the Patient Health Questionnaire (PHQ-9) (Kroenke et al., 2001), verbal intellectual function was assessed with the National Adult Reading Test (NART) (Nelson, 1991) and cognitive impairment with the Mini Mental State Exam (MMSE) (Folstein et al., 1975). One participant was excluded after assessment of their MMSE score (MMSE = 26).

**Table 1.** Participant Demographics. This table lists the demographics (age, years of education, sex) of the participants that took part in this study, and splits male (M) and female (F) data by risk factor group. FH = family history of dementia, APOE4 = Apolipoprotein ε4, and waist-hip ratio. Mean age and years of education, accurate to 2 decimal places, are quoted with standard deviations reported in brackets (σ).

|  |  |  |  |  |
| --- | --- | --- | --- | --- |
| | Mean ($\sigma$) | | | |
| Age | 55.76 (8.22), Range: 38-71 |  |  |  |
| Males | 71/165 |  |  |  |
| Years of Education | 16.55 (3.32), Range: 9.5-26 |  |  |  |
| FH of dementia | 59/163 |  |  |  |
| APOE4 carriers | 64/164 |  |  |  |
| Waist Hip Ratio – Obese | 102/165 |  |  |  |
| Demographics broken down by risk factor |  |  |  |  |
| | N (M) | N (F) | Mean age, M ($\sigma$) | Mean age, F ($\sigma$) |
| No FH, No APOE4, Healthy weight | 4 | 16 | 53.75 (4.03) | 53.69 (8.68) |
| FH, No APOE4, Healthy weight | 0 | 13 | - | 53.85 (6.87) |
| No FH, APOE4, Healthy weight | 3 | 16 | 49.00 (7.21) | 52.88 (9.64) |
| No FH, No APOE4, Obese | 18 | 21 | 56.17 (8.74) | 56.05 (7.68) |
| FH, APOE4, Healthy weight | 4 | 6 | 59.00 (2.45) | 59.83 (5.19) |
| No FH, APOE4, Obese | 20 | 6 | 54.00 (9.61) | 58.67 (6.56) |
| FH, No APOE4, Obese | 17 | 10 | 58.71 (7.71) | 56.60 (9.35) |
| FH, APOE4, Obese | 5 | 3 | 57.00 (8.22) | 62.00 (7.94) |

### 1.2 Assessment of risk factors

Participants gave saliva samples with the Genotek Oragene-DNA kit (OG-500) for *APOE* genotyping. *APOE* genotypes ε2, ε3 and ε4 were determined by TaqMan genotyping of single nucleotide polymorphism (SNP) rs7412 and KASP genotyping of SNP rs429358 (Metzler-Baddeley, Mole, Leonaviciute, et al., 2019). Genotyping was successful for 164 of the 165 participants. In addition, 163 participants provided information about their family history of dementia, i.e. whether a first-grade relative was affected by AD, vascular dementia or any other type of dementia. We also obtained the number of years spent in education for 164 participants, to include as a covariate in this analysis (Table 1).

Participants’ waist and hip circumferences were measured to calculate the waist-hip-ratio (WHR). Central obesity was defined as a WHR ≥ 0.9 for men and ≥ 0.85 for women (Table 1). Other metabolic risk factors were self-reported in a medical history questionnaire (see for details Mole et al., 2020 Neurobiology of Aging) but were not included in the present analysis.

### 1.3 MRI data acquisition

MRI data were collected on a 3T MAGNETOM Prisma clinical scanner (Siemens Healthcare, Erlangen, Germany) (Coad et al., 2020; Metzler-Baddeley, Mole, Leonaviciute, et al., 2019; Metzler-Baddeley, Mole, Sims, et al., 2019; Mole, Fasano, Evans, Sims, Hamilton, et al., 2020) at the Cardiff University Brain Research Imaging Centre (CUBRIC). A 3D magnetization-prepared rapid gradient-echo (MP-RAGE) sequence was used to acquire T_1_-weighted anatomical images with the following parameters: 256×256 acquisition matrix, TR = 2300 ms, TE = 3.06 ms, TI = 850 ms, flip angle θ = 9°, 176 slices, 1 mm slice thickness, 1×1×1 mm isotropic resolution, FOV = 256 mm and acquisition time of ∼ 6 min.

Diffusion-weighted MR images were acquired with High Angular Resolution Diffusion Imaging (HARDI) (Tuch et al., 2002) using a spin-echo echo-planar dual shell HARDI sequence with diffusion encoded along 90 isotropically distributed orientations (Jones et al., 1999) (30 directions at b-value = 1200 s/mm^2^, 60 directions at b-value = 2400 s/mm^2^) as well as six non-diffusion weighted images with dynamic field correction using the following parameters: TR = 9400ms, TE = 67ms, 80 slices, 2 mm slice thickness, 2×2×2 mm voxel, FOV = 256×256×160 mm, GRAPPA acceleration factor = 2, acquisition time of ∼15 min.

### 1.4 HARDI data processing and whole brain tractography

Diffusion-weighted imaging data processing has been previously detailed in Coad et al., 2020; Metzler-baddeley, Mole, Leonaviciute, et al., 2019; Metzler-baddeley, Mole, Sims, et al., 2019; Mole et al., 2020. In brief, dual-shell data were split and b = 1200 and 2400 s/mm^2^ data were corrected separately for distortions induced by the diffusion-weighted gradients and motion artefacts in ExploreDTI (v4.8.3) (Leemans et al., 2009). EPI-induced geometrical distortions were corrected by registering the diffusion-weighted image volumes to the T_1_-weighted images (Irfanoglu et al., 2012).

Outliers in the diffusion data were identified with the RESDORE algorithm (Parker, 2014). Whole brain tractography was performed with the damped Richardson-Lucy algorithm (dRL) (Dell’Acqua et al., 2010) on the 60 direction, b = 2400 s/mm^2^ HARDI data for each dataset in single-subject space using in-house software (Parker, 2014) coded in MATLAB (the MathWorks, Natick, MA). Fibre tracts were reconstructed by estimating the dRL fibre orientation density functions (fODFs) at the centre of each image voxel with seed points positioned at the vertices of a 2×2×2 mm grid superimposed over the image. At each seed point the tracking algorithm interpolated local fODF estimates and then propagated 0.5 mm along orientations of each fODF lobe above a threshold of a peak amplitude of 0.05. Individual streamlines were then propagated by interpolating the fODF at their new location and by propagating 0.5 mm along the minimally subtending fODF peak. This process was repeated until the minimally subtending peak magnitude fell below 0.05 or the change of direction exceeded an angle of 45°. Tracking was subsequently repeated in the opposite direction from the initial seed point. Streamlines with lengths outside a range of 10 mm to 500 mm were removed.

### 1.5 Generating integrated weighted structural brain networks: whole-brain analysis

Whole brain tractography maps were used in ExploreDTI v4.8.6 (Leemans et al., 2009), to create connectivity matrices that describe the structural connectome mathematically. Network nodes were defined according to the automated anatomical labelling (AAL) atlas (Tzourio-Mazoyer et al., 2002) using the 90 cortical and subcortical areas of the cerebrum. The edges of the networks were the tractography-reconstructed tracts: all edges between brain areas not connected by tracts were therefore equal to zero. This process resulted in 16 90×90 connectivity matrices, the edges of each quantifying if there was a tract or not, number of tracts between two nodes, percentage of tracts between two nodes, average tract length, Euclidean distance, density of tracts, tract volume, mean diffusivity, axial diffusivity, radial diffusivity, fractional anisotropy, second and third eigenvalue of the diffusion tensor, linear anisotropy, planar anisotropy and spherical anisotropy.

The above-mentioned metrics are chosen because they could reflect the signal transport and integration abilities of the structural connectome. However, it is not clear yet to what extent they achieve that (Messaritaki et al., 2021). Additionally, the strength of the structural connectivity between brain areas depends on the metric used to weight the network edges. As a result, the network measures derived via the graph theoretical analysis depend on the connectivity matrix used – i.e. which of the above metrics we chose as an edge weight. We have recently shown that this ambiguity can be solved by linearly combining nine normalised metrics (number of tracts, percentage of tracts, average tract length, Euclidean distance, density, tract volume, mean diffusivity, radial diffusivity and fractional anisotropy) into a single graph (Dimitriadis, Drakesmith, et al., 2017) and thresholding the subsequent graphs using an orthogonal minimal spanning trees scheme (Dimitriadis, Antonakakis, et al., 2017). This protocol creates connectivity matrices that combine the information from the included metrics in a data driven manner, so that the maximum information from all metrics is retained in the final graph; these are termed integrated graphs. The thresholding step can be applied in dense matrices, resulting in a topographically filtered integrated weighted structural brain network. The network and nodal reliability of such integrated graphs was improved beyond that of the 9 individual metrics (Dimitriadis, Drakesmith, et al., 2017). In addition, they were shown to have very good discrimination capability in a binary classification problem (Dimitriadis, Drakesmith, et al., 2017), and to exhibit good scan-rescan reliability (Messaritaki et al., 2019b, 2019a). A recent study demonstrated that community partitions and provincial hubs are highly reproducible in a test-retest study when structural brain networks were constructed with the integrated approach (Dimitriadis et al., 2020). For those reasons we created integrated weighted brain networks instead of pursuing a single-metric structural connectivity matrix.

In order to reduce the number of false positives possibly resulting from the tractography, we set to zero all edges in the structural connectivity matrices that corresponded to tracts with fewer than 5 streamlines (excluding Euclidean distance as this is a biological metric and has a value regardless of the number of streamlines). All subsequent analyses were performed on these thresholded connectivity matrices (Figure 1).

**Figure 1.**
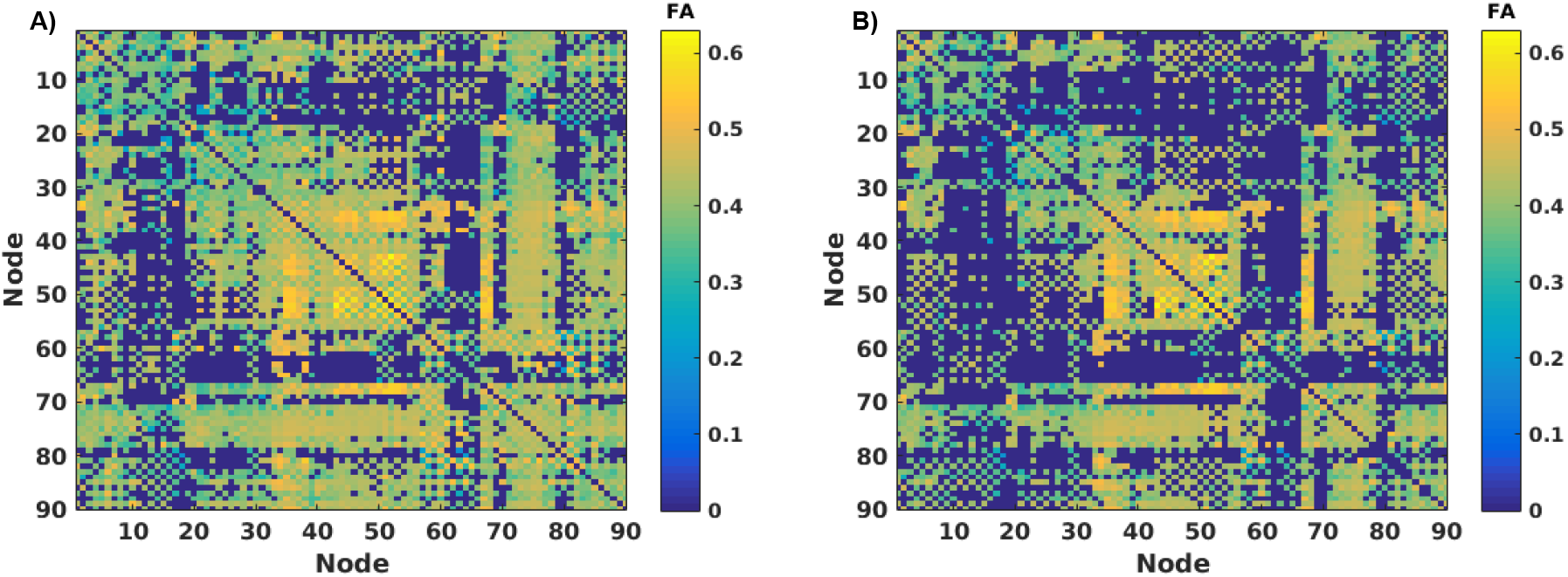
An example of the conservative threshold added to all dMRI connectivity matrices. **A)** Fractional anisotropy (FA) connectivity matrix for one participant before thresholding. **B)** After a conservative threshold of 5 streamlines was applied to FA for the same participant.

In order to decide which metrics to combine into the integrated weighted structural brain network, we calculated the intercorrelation coefficients (Corrcoef, MATLAB R2015a) between the number of tracts (NS), percentage of tracts (PS), average tract length (ATL), Euclidean distance (ED), density of streamlines (SLD), tract volume (TV), mean diffusivity (MD), radial diffusivity (RD), axial diffusivity (AxD) and fractional anisotropy (FA), see Table 2. In addition, we performed a multicollinearity test (Collintest, MATLAB R2015a) in an endeavour to eliminate metrics representing redundant information within our integrated graphs. After excluding highly correlated and multi-collinear metrics, the remaining metrics were integrated into a single graph via a linear graph-distance combination (Dimitriadis, Drakesmith, et al., 2017)^1^.

**Table 2.** Abbreviations used for the dMRI metrics. This table defines the abbreviations used throughout the manuscript for each of the diffusion-weighted magnetic resonance imaging (dMRI) metrics.

| **Name of dMRI metric** | **Abbreviation** |
| --- | --- |
| Number of tracts | NS |
| Percentage of tracts, | PS |
| Average tract length | ATL |
| Euclidean distance | ED |
| Streamline density | SLD |
| Tract volume | TV |
| Mean diffusivity | MD |
| Radial diffusivity | RD |
| Axial diffusivity | AxD |
| Fractional anisotropy | FA |

### 1.6 Calculating network measures from integrated graphs

The resulting graphs were weighted and undirected. Using the MATLAB Brain Connectivity Toolbox (Rubinov & Sporns, 2010) we calculated the following metrics:

- Clustering coefficient: A measure of how interconnected nodes are (averaged across all nodes)
- Characteristic path length: The average minimum number of connections to span two nodes
- Eccentricity: Maximum shortest distance between one node and all others (averaged across all nodes)
- Radius: Minimum eccentricity
- Diameter: Maximum eccentricity
- Global efficiency: Inverse of the characteristic path length^2^

Network measures were examined for multicollinearity using Belsley collinearity diagnostics (Collintest, MATLAB R2015a) to ensure only unique predictors were included in our analysis. The remaining network measures were analysed using multivariate general linear models described below. We were also interested in identifying potential interactions between our risk factors.

### 1.7 Sub-network analysis

As AD preferentially impacts the DMN we repeated the analysis for this sub-network by adapting the AAL atlas (Tzourio-Mazoyer et al., 2002) based on the data from Power et al., 2011. The DMN graphs were comprised of 22 nodes from each hemisphere encompassing the frontal, temporal, parietal lobes including the precuneus, cingulate gyrus and hippocampus (Figure 2). To investigate if any changes were specific to the DMN we analysed a separate control sub-network – the visual system (Wang et al., 2012) by adjusting the regions of interest specified in Power et al., 2011. The resulting integrated weighted structural brain networks were composed of 16 nodes from the left and right hemispheres: inferior temporal gyrus, fusiform gyrus, superior/middle/inferior occipital gyrus, lingual gyrus, cuneus and calcarine fissure and surrounding cortex (Figure 2).

**Figure 2.**
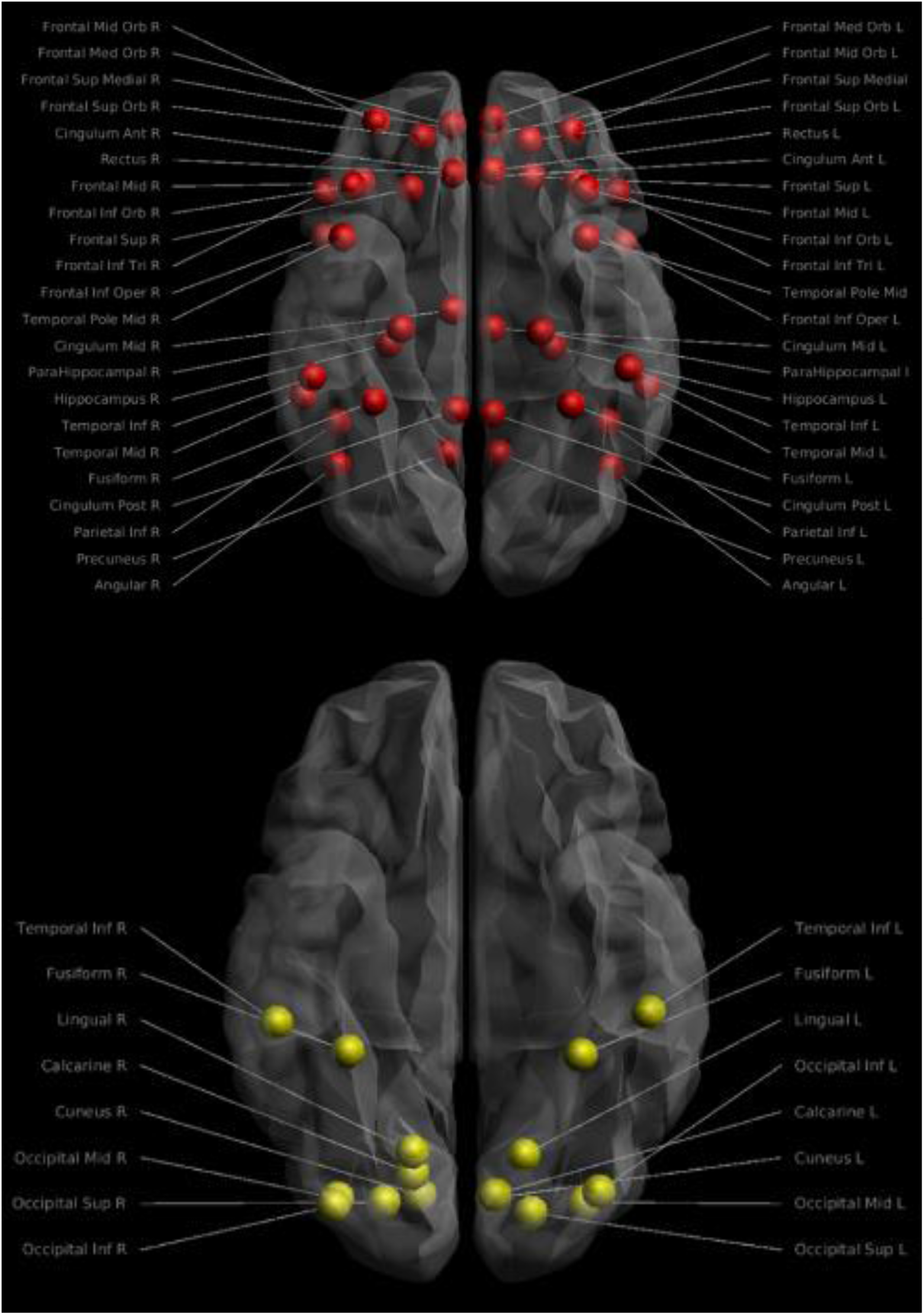
Nodes included in the subnetwork analysis for the default mode network (DMN) and visual system. The top figure shows the 44 nodes included in the DMN adapted from Power et al., 2011 whereas the bottom picture shows the 16 nodes included in the visual network adapted from Power et al., 2011. Images were created using ExploreDTI v4.8.6.

### 1.8 Hub analysis

Hubs are nodes of a network that are highly connected to other nodes and act as bridges that facilitate the transfer of signals in the brain, contributing to its integration abilities (van den Heuvel & Sporns, 2013). Crucially, hubs appear to play a role in AD (Buckner et al., 2009). We split the cohort into risk factor groups – positive (N = 59) vs. negative family history (N = 104), APOE4 carriers (N = 64) vs. non-carriers (N = 100), centrally obese (N = 102) vs. healthy weight (N = 63) – to explore whether hubs changed as a function of risk factor profile in healthy individuals. Hubs were identified across the whole-brain for each participant by ranking nodal betweenness centrality and strength, where higher scores indicate hubs. In addition, nodal local efficiency and clustering coefficients were ranked, with smaller values indicating hubs. A node was defined as a hub when it was in the top 20% for global measures and the lowest 20% for local measures. Using replicator dynamics (Dimitriadis et al., 2010; Neumann et al., 2005), hubs which were consistently present across the individual risk factor cohorts were determined^3^. This analysis was then repeated using data from the DMN and visual sub-networks to identify internally important nodes.

### 1.9 Statistical analyses

In SPSS v26 (IBM Corp., 2019), we performed multivariate general linear models with factors of APOE4 carrier/non-carrier, family history of dementia/no family history and waist-hip ratio obese/healthy on dependent variables; mean clustering coefficient, characteristic path length, eccentricity, global efficiency, diameter and radius. The analyses were adjusted for covariates: age, years of education and sex. To ensure assumptions were met, normality of residuals was tested using Kolmogorov-Smirnov tests. We adopted Belsley collinearity diagnostics (MATLAB R2015a) to assess multicollinearity effects between the estimated network metrics.

## 2. Results

### 2.1 Inclusion of metrics into integrated networks using correlation and collinearity tests

A multicollinearity test was performed on the 10 variables with a default cut-off of 30 for the condition index and 0.5 for proportion of variance-decomposition. This analysis revealed multicollinearity between AxD, MD and RD (Table 3). Correlation coefficients (Table 4) were calculated between all 10 connectivity metrics. We used a cut off of R > 0.6 to flag strong correlations to investigate further. PS, NS and TV were highly inter-correlated, and for that reason we only included NS in our analysis. AxD, MD and RD exhibit multicollinearity and both AxD and RD correlated strongly with FA (R = 0.6036, p-value < 10^−8^ and R = -0.6721, p-value < 10^−8^, respectively) – thus these two metrics were excluded. This resulted in a final inclusion of ATL, SLD, FA, ED, MD and NS. We re-ran the correlation and multicollinearity analysis on these metrics and confirmed no strong correlations (Table 4) or multicollinearity. These six metrics were then combined into a single graph (Figure 3) with an algorithm introduced in our previous study (Dimitriadis, Drakesmith, et al., 2017).

**Table 3.** Belsley collinearity diagnostics results for dMRI connectivity matrices. Belsley collinearity diagnostics run across the diffusion-weighted magnetic resonance imaging (dMRI) metrics demonstrated multicollinearity between AxD, MD and RD. The highlighted cells identify metrics which meet our exclusion criteria, condition index > 30 and variance decomposition > 0.5. sValue = Singular values, CondIdx = Condition Index. Abbreviations of the dMRI metrics are defined in Table 2.

| sValue | Condlx | AxD | ATL | ED | FA | MD | NS | PS | RD | SLD | TV |
| --- | --- | --- | --- | --- | --- | --- | --- | --- | --- | --- | --- |
| 2.7153 | 1 | 0 | 0.001 | 0.0013 | 0.0001 | 0 | 0.0004 | 0.0005 | 0 | 0.0029 | 0.001 |
| 1.3103 | 2.0722 | 0 | 0.0021 | 0.0047 | 0.0001 | 0 | 0.0074 | 0.0104 | 0 | 0 | 0.0082 |
| 0.7989 | 3.399 | 0 | 0.0073 | 0.0065 | 0 | 0 | 0.0002 | 0.0005 | 0 | 0.4447 | 0.0034 |
| 0.3047 | 8.91 | 0 | 0.0299 | 0.3087 | 0.0001 | 0 | 0.0155 | 0.1257 | 0 | 0.0001 | 0.3775 |
| 0.2837 | 9.5698 | 0 | 0.0521 | 0.3839 | 0.0063 | 0 | 0.0018 | 0.043 | 0 | 0.3094 | 0.1527 |
| 0.2258 | 12.0263 | 0 | 0.7593 | 0.1808 | 0 | 0 | 0.0041 | 0.1228 | 0 | 0.1427 | 0.1425 |
| 0.1587 | 17.1051 | 0 | 0.0497 | 0.0493 | 0.0007 | 0 | 0.969 | 0.693 | 0 | 0.0078 | 0.3067 |
| 0.1433 | 18.9544 | 0 | 0.085 | 0.0034 | 0.2084 | 0 | 0.0016 | 0.0037 | 0 | 0.0732 | 0.0074 |
| 0.0414 | 65.5353 | 0 | 0.0135 | 0.0614 | 0.7839 | 0 | 0 | 0.0003 | 0 | 0.0192 | 0.0005 |
| 0 | 2.26E+14 | 1 | 0 | 0.0001 | 0.0004 | 1 | 0 | 0 | 1 | 0 | 0 |

**Table 4.**
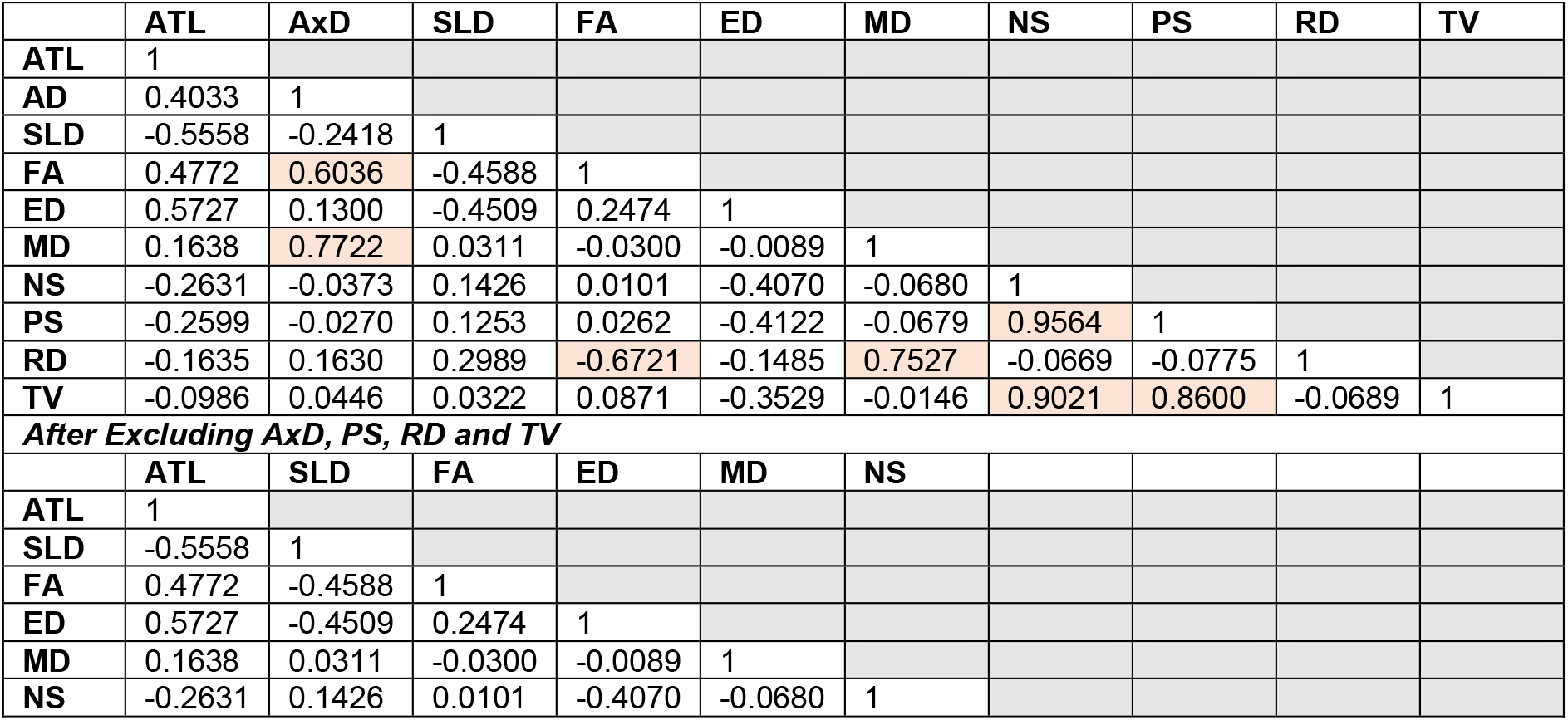
Correlation coefficients (R) as determined by MATLAB (corrcoef) between the individual connectivity metrics (abbreviations defined in table 2). Highlighted cells identify inter-correlations with an R > 0.6. The lower half of the table shows reduced inter-correlation coefficients after the analysis has been re-run with AxD, PS, RD and TV excluded.

|  | ATL | AxD | SLD | FA | ED | MD | NS | PS | RD | TV |
| --- | --- | --- | --- | --- | --- | --- | --- | --- | --- | --- |
| ATL | 1 |  |  |  |  |  |  |  |  |  |
| AD | 0.4033 | 1 |  |  |  |  |  |  |  |  |
| SLD | -0.5558 | -0.2418 | 1 |  |  |  |  |  |  |  |
| FA | 0.4772 | 0.6036 | -0.4588 | 1 |  |  |  |  |  |  |
| ED | 0.5727 | 0.1300 | -0.4509 | 0.2474 | 1 |  |  |  |  |  |
| MD | 0.1638 | 0.7722 | 0.0311 | -0.0300 | -0.0089 | 1 |  |  |  |  |
| NS | -0.2631 | -0.0373 | 0.1426 | 0.0101 | -0.4070 | -0.0680 | 1 |  |  |  |
| PS | -0.2599 | -0.0270 | 0.1253 | 0.0262 | -0.4122 | -0.0679 | 0.9564 | 1 |  |  |
| RD | -0.1635 | 0.1630 | 0.2989 | -0.6721 | -0.1485 | 0.7527 | -0.0669 | -0.0775 | 1 |  |
| TV | -0.0986 | 0.0446 | 0.0322 | 0.0871 | -0.3529 | -0.0146 | 0.9021 | 0.8600 | -0.0689 | 1 |
| *After Excluding AxD, PS, RD and TV* |  |  |  |  |  |  |  |  |  |  |
|  | ATL | SLD | FA | ED | MD | NS |  |  |  |  |
| ATL | 1 |  |  |  |  |  |  |  |  |  |
| SLD | -0.5558 | 1 |  |  |  |  |  |  |  |  |
| FA | 0.4772 | -0.4588 | 1 |  |  |  |  |  |  |  |
| ED | 0.5727 | -0.4509 | 0.2474 | 1 |  |  |  |  |  |  |
| MD | 0.1638 | 0.0311 | -0.0300 | -0.0089 | 1 |  |  |  |  |  |
| NS | -0.2631 | 0.1426 | 0.0101 | -0.4070 | -0.0680 | 1 |  |  |  |  |

**Figure 3.**
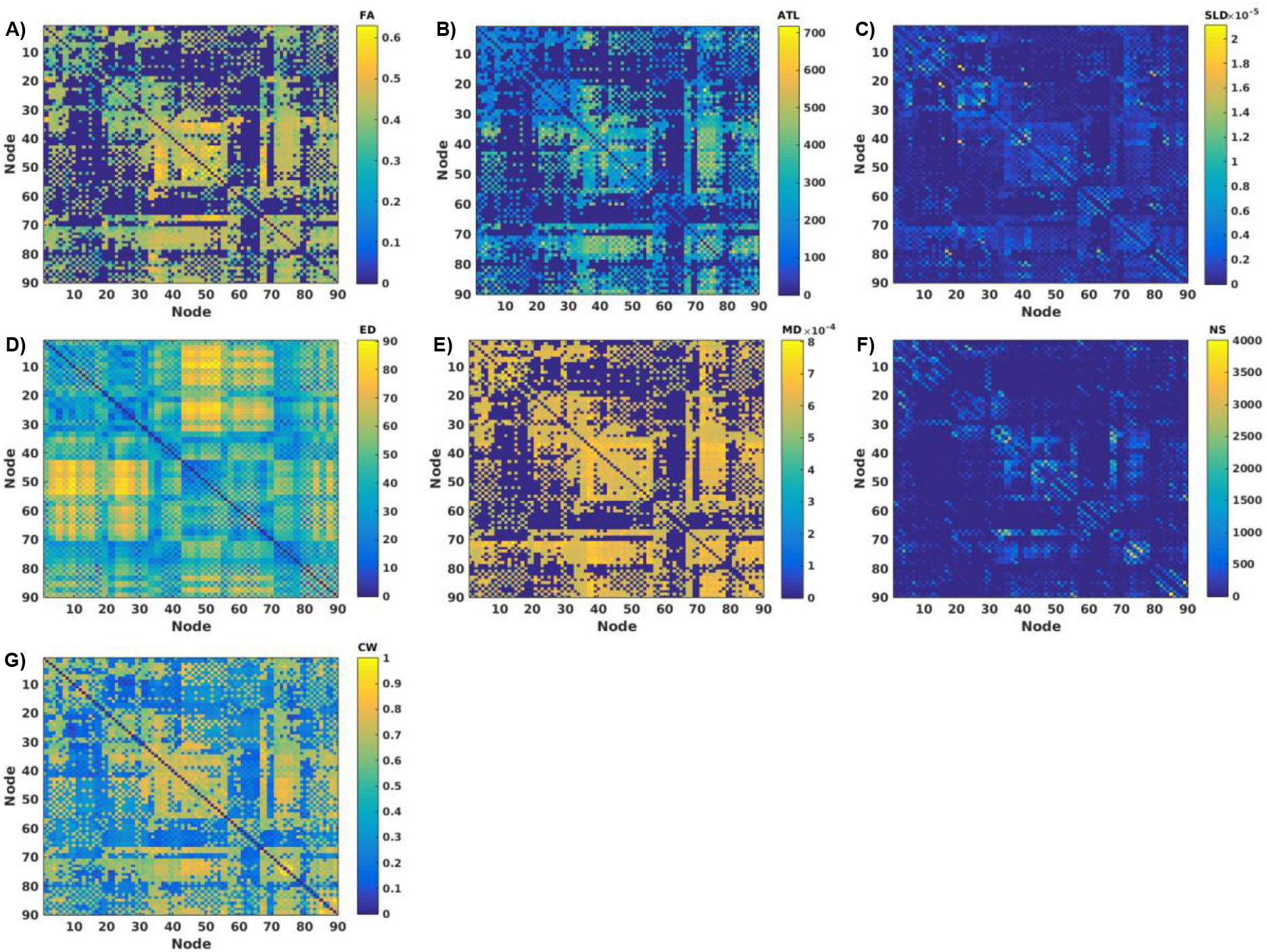
An example of the six Individual network measures that were combined into an integrated weighted structural brain network for one participant. **A)** FA, fractional anisotropy, **B)** ATL, average tract length **C)** SLD, streamline density, **D)** ED, Euclidean distance, **E)** MD, mean diffusivity **F)** NS, number of streamlines were combined into a **G)** integrated weighted structural brain network. CW = connectivity weight.

### 2.2. Exclusion of mean eccentricity from further analyses

Belsley collinearity diagnostics applied over the adopted set of network metrics flagged multicollinearity between diameter and mean eccentricity using whole-brain network measures (Table 5). We therefore excluded the eccentricity from further analyses and kept the diameter, which in combination to radius inform us about the lower (radius) and upper limits (diameter) of eccentricity.

**Table 5.** Belsley collinearity diagnostics results - network measures. This table demonstrates multicollinearity between whole-brain diameter and mean eccentricity when assessed with Belsley collinearity diagnostics. The highlighted cells meet exclusion criteria. sValue = Singular values, CondIdx = Condition Index.

| sValue | Condlx | Diameter | Efficiency | Lambda | Radius | Clustering coefficient | Eccentricity |
| --- | --- | --- | --- | --- | --- | --- | --- |
| 2.4172 | 1.0000 | 0.0001 | 0.0003 | 0.0001 | 0.0000 | 0.0004 | 0.0000 |
| 0.3748 | 6.4489 | 0.0006 | 0.0257 | 0.0038 | 0.0010 | 0.0374 | 0.0003 |
| 0.0890 | 27.1550 | 0.0440 | 0.3625 | 0.1359 | 0.0011 | 0.5189 | 0.0002 |
| 0.0812 | 29.7691 | 0.0640 | 0.5520 | 0.1607 | 0.0000 | 0.3796 | 0.0018 |
| 0.0395 | 61.2534 | 0.2325 | 0.0107 | 0.2996 | 0.7250 | 0.0146 | 0.0004 |
| 0.0205 | 118.0614 | 0.6589 | 0.0487 | 0.4000 | 0.2729 | 0.0492 | 0.9972 |

### 2.3 Whole-brain analysis

Kolmogorov-Smirnov tests for normality revealed non-Gaussian distributions for all network measures (p-value < 0.05, Table 6, Figure S1). To alleviate this, diameter, characteristic path length and radius were log transformed to reduce positive skew, and efficiency and clustering coefficients were squared to reduce negative skew. After data cleaning, the diameter, characteristic path length, and radius mimicked normal distributions when assessed by Kolmogorov-Smirnov tests (p-value > 0.05), but efficiency and clustering coefficients were non-normal. Despite the latter, the analysis was continued as the graphs, when inspected by eye, appeared improved beyond the original in regard to skew (Figure S1), and therefore we decided that the data complied with general linear model assumptions despite formally failing the tests. One extreme outlier was present in the diameter data after transformation (defined as > 3 x interquartile range) thus we excluded this participant from further whole-brain analyses. Four participants had missing data thus the final analysis had a sample size of 160. Omnibus multivariate analyses revealed no significant main or interaction effects (p-value > 0.05, Table 7), suggesting that there were no differences in whole-brain network measures between individuals who carry APOE4 vs. non-carriers, have a FH vs. no FH, obese vs. healthy weight.

**Table 6.** Kolmogorov-Smirnov test results for the whole-brain, DMN and visual system. Lack of normality of standardized residuals assessed with Kolmogorov-Smirnov tests of the whole-brain, DMN (default mode network) and visual system. The lower part of the table demonstrates how the normality of the metrics are improved after data cleaning (removing outliers and transforming the data). DF = degrees of freedom.

| **Before Data Cleaning** |  |  |  |  |
| --- | --- | --- | --- | --- |
|  | **Standardised Residual** | **Statistic** | **DF** | **p-value** |
| **Whole** | Diameter | 0.078 | 161 | 0.019 |
|  | Global efficiency | 0.114 | 161 | 0.000 |
|  | Characteristic path length | 0.082 | 161 | 0.010 |
|  | Radius | 0.083 | 161 | 0.008 |
|  | Clustering coefficient | 0.115 | 161 | 0.000 |
| **DMN** | Diameter | 0.118 | 161 | 0.000 |
|  | Global efficiency | 0.200 | 161 | 0.000 |
|  | Characteristic path length | 0.158 | 161 | 0.000 |
|  | Radius | 0.120 | 161 | 0.000 |
|  | Clustering coefficient | 0.146 | 161 | 0.000 |
| **Visual** | Diameter | 0.128 | 161 | 0.000 |
|  | Global efficiency | 0.128 | 161 | 0.000 |
|  | Characteristic path length | 0.124 | 161 | 0.000 |
|  | Radius | 0.135 | 161 | 0.000 |
|  | Clustering coefficient | 0.086 | 161 | 0.006 |
| **After Data Cleaning** |  |  |  |  |
|  | **Standardised Residual** | **Statistic** | **DF** | **p-value** |
| **Whole** | Logged diameter | 0.067 | 160 | 0.080 |
|  | Squared global efficiency | 0.091 | 160 | 0.003 |
|  | Logged characteristic path length | 0.047 | 160 | 0.200 |
|  | Logged radius | 0.053 | 160 | 0.200 |
|  | Squared clustering coefficient | 0.083 | 160 | 0.009 |
| **DMN** | Logged diameter | 0.059 | 151 | 0.200 |
|  | Squared global efficiency | 0.151 | 151 | 0.000 |
|  | Logged characteristic path length | 0.084 | 151 | 0.010 |
|  | Logged Radius | 0.060 | 151 | 0.200 |
|  | Clustering coefficient | 0.123 | 151 | 0.000 |
| **Visual** | Logged diameter | 0.091 | 161 | 0.003 |
|  | Squared global efficiency | 0.108 | 161 | 0.000 |
|  | Logged characteristic path length | 0.082 | 161 | 0.009 |
|  | Logged radius | 0.088 | 161 | 0.004 |
|  | Squared clustering coefficient | 0.060 | 161 | 0.200 |

**Table 7.** Multivariate results. There were no significant differences in network measures as a function of risk factors: family history of dementia (FH), Apolipoprotein ε4 genotype (APOE4) and waist-hip ratio (WHR) across the whole-brain, default mode network or a control sub-network (visual system). F = F statistic, DF = degrees of freedom.

| **Whole-brain analysis** |  |  |  |
| --- | --- | --- | --- |
| **Effect** | **F** | **DF** | **p-value** |
| Intercept | 48.648 | 5, 145 | 0.000 |
| Sex | 0.841 | 5, 145 | 0.523 |
| Age | 1.325 | 5, 145 | 0.257 |
| Years of Education | 1.904 | 5, 145 | 0.097 |
| FH | 1.307 | 5, 145 | 0.264 |
| APOE4 | 0.351 | 5, 145 | 0.881 |
| WHR | 0.981 | 5, 145 | 0.432 |
| FH X APOE4 | 1.019 | 5, 145 | 0.409 |
| FH X WHR | 0.532 | 5, 145 | 0.752 |
| APOE4 X WHR | 0.533 | 5, 145 | 0.751 |
| FH X APOE4 X WHR | 1.666 | 5, 145 | 0.147 |
| **Default mode network analysis** |  |  |  |
| Intercept | 84.361 | 5, 136 | 0.000 |
| Sex | 1.315 | 5, 136 | 0.261 |
| Age | 1.867 | 5, 136 | 0.104 |
| Years of Education | 1.010 | 5, 136 | 0.414 |
| FH | 1.523 | 5, 136 | 0.187 |
| APOE4 | 0.924 | 5, 136 | 0.567 |
| WHR | 0.201 | 5, 136 | 0.961 |
| FH X APOE4 | 0.242 | 5, 136 | 0.242 |
| FH X WHR | 0.733 | 5, 136 | 0.733 |
| APOE4 X WHR | 0.940 | 5, 136 | 0.940 |
| FH X APOE4 X WHR | 0.444 | 5, 136 | 0.444 |
| **Visual system analysis** |  |  |  |
| Intercept | 73.555 | 5, 146 | 0.000 |
| Sex | 0.534 | 5, 146 | 0.750 |
| Age | 1.989 | 5, 146 | 0.084 |
| Years of Education | 1.314 | 5, 146 | 0.261 |
| FH | 0.351 | 5, 146 | 0.881 |
| APOE4 | 0.901 | 5, 146 | 0.482 |
| WHR | 1.179 | 5, 146 | 0.322 |
| FH X APOE4 | 0.284 | 5, 146 | 0.921 |
| FH X WHR | 0.986 | 5, 146 | 0.429 |
| APOE4 X WHR | 1.365 | 5, 146 | 0.241 |
| FH X APOE4 X WHR | 2.089 | 5, 146 | 0.070 |

### 2.4 Sub-network analyses

We then investigated whether any individual differences were occurring at a sub-network level.

#### 2.4.1 Default Mode Network analysis

Kolmogorov-Smirnov tests revealed non-normality for all network measures calculated from the DMN integrated graphs (Table 6, Figure S2). To correct for this, the diameter, characteristic path length and radius were log transformed and the efficiency was squared. Despite being non-Gaussian as determined by Kolmogorov-Smirnov tests, the distribution of residuals for mean clustering coefficients was not heavily skewed when assessed by eye. We identified nine extreme outliers (> 3 x interquartile range) within efficiency data and one within clustering coefficients thus these were removed from the analysis. After data cleaning, the characteristic path length, efficiency and clustering coefficients were not formally normal when reassessed with Kolmogorov-Smirnov tests, however the analysis was continued (Figure S2) as not to lose value in our raw data as a result another round of data cleaning. Multivariate analyses revealed no significant effects (N = 151, p-value > 0.05, Table 7) suggesting that there are no differences in default mode network measures as a result of risk-factor profile.

#### 2.4.2 Visual network analysis

Kolmogorov-Smirnov tests revealed non-normality for all six network measures for the visual system (Table 6, Figure S3). Following the same process as before, the diameter, characteristic path length and radius were log transformed and the efficiency and clustering coefficients were squared. No outliers were identified in the transformed metrics. Diameter, characteristic path length, radius and efficiency were non-normal, however the analysis was continued (Figure S3), with a sample size of 161, as the distributions were improved beyond the untransformed metrics to a point which we believe meets the underlying assumptions of the analysis. The general linear model (N = 161) revealed no significant multivariate effects (Table 7).

### 2.5 Analysis of network hubs in the whole-brain

Replicator dynamics identified hubs consistent across the individual risk factor groups. Individuals with no FH (N = 104) had hubs located in the left and right rolandic operculum, right inferior parietal gyrus, left angular gyrus and right Heschl’s gyrus, whereas individuals with a positive FH (N = 59) had hubs at the right rolandic operculum, left inferior frontal gyrus opercular part, left and right paracentral lobule and the right Heschl’s gyrus (Figure 4). Individuals who were of healthy weight (N = 63), and thus considered at less risk of developing AD, had hubs within the left inferior frontal gyrus opercular part, right rolandic operculum, right inferior parietal gyrus and right Heschl’s gyrus, whereas individuals who were obese (N = 102) had hubs within the right rolandic operculum, right paracentral lobule and both left and right Heschl’s gyrus (Figure 4). Participants who were negative for the APOE4 allele (and thus considered low-risk), had hubs in the left inferior frontal gyrus opercular part, right rolandic operculum, right precuneus and right Heschl’s gyrus (N = 100) whereas APOE4 positive individuals (N = 64) had hubs in the right rolandic operculum, right inferior parietal gyrus, left angular gyrus, right paracentral lobule and right Heschl’s gyrus (Figure 4).

**Figure 4.**
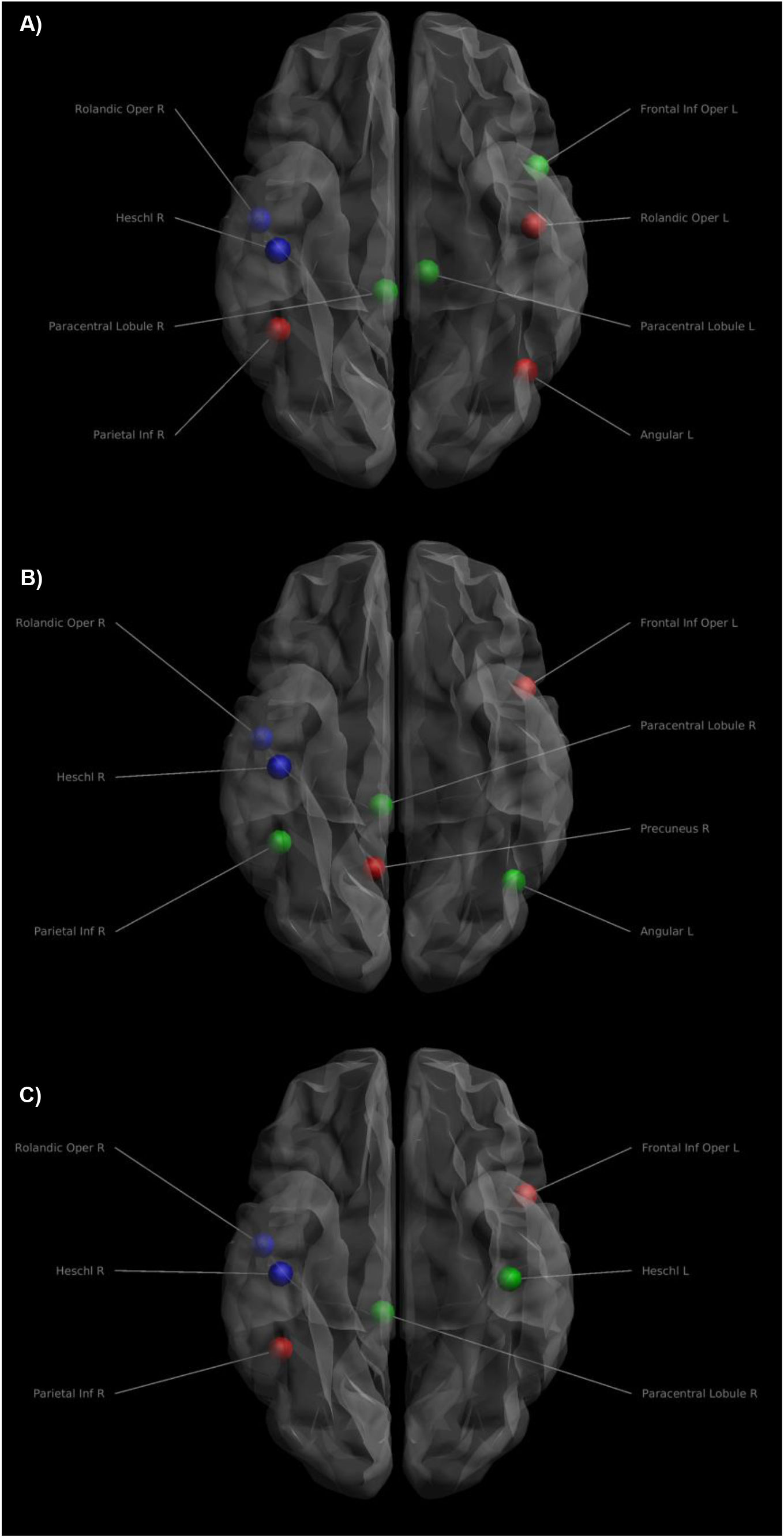
Nodes identified as hubs change dependent on risk factor profile. This figure shows the changes in nodes defined as hub regions, when you transition from a low-risk group to a high-risk group. **A)** Comparing individuals without a family history of dementia (FH) to those with a positive FH indicates that 2 hubs remain unchanged (blue), whereas 3 are gained (green) and 3 are lost (red). **B)** Comparing APOE4 non-carriers to carriers results in a gain of 3 hubs (green), loss of 2 hubs (red) but leaves 2 hubs unchanged (blue). **C)** In comparison to healthy individuals, obese participants gained 2 hubs (green), lost 2 hubs (red) and 2 hubs remains (blue).

To summarise the above described pattern, the right rolandic operculum and Heschl’s gyrus remain present as hubs regardless of risk factor. In contrast, at risk individuals (obese, positive FH and APOE4 carriers) consistently have a hub in the right paracentral lobule which is absent in their respective low-risk group (Table 8, Figure 4).

**Table 8.**
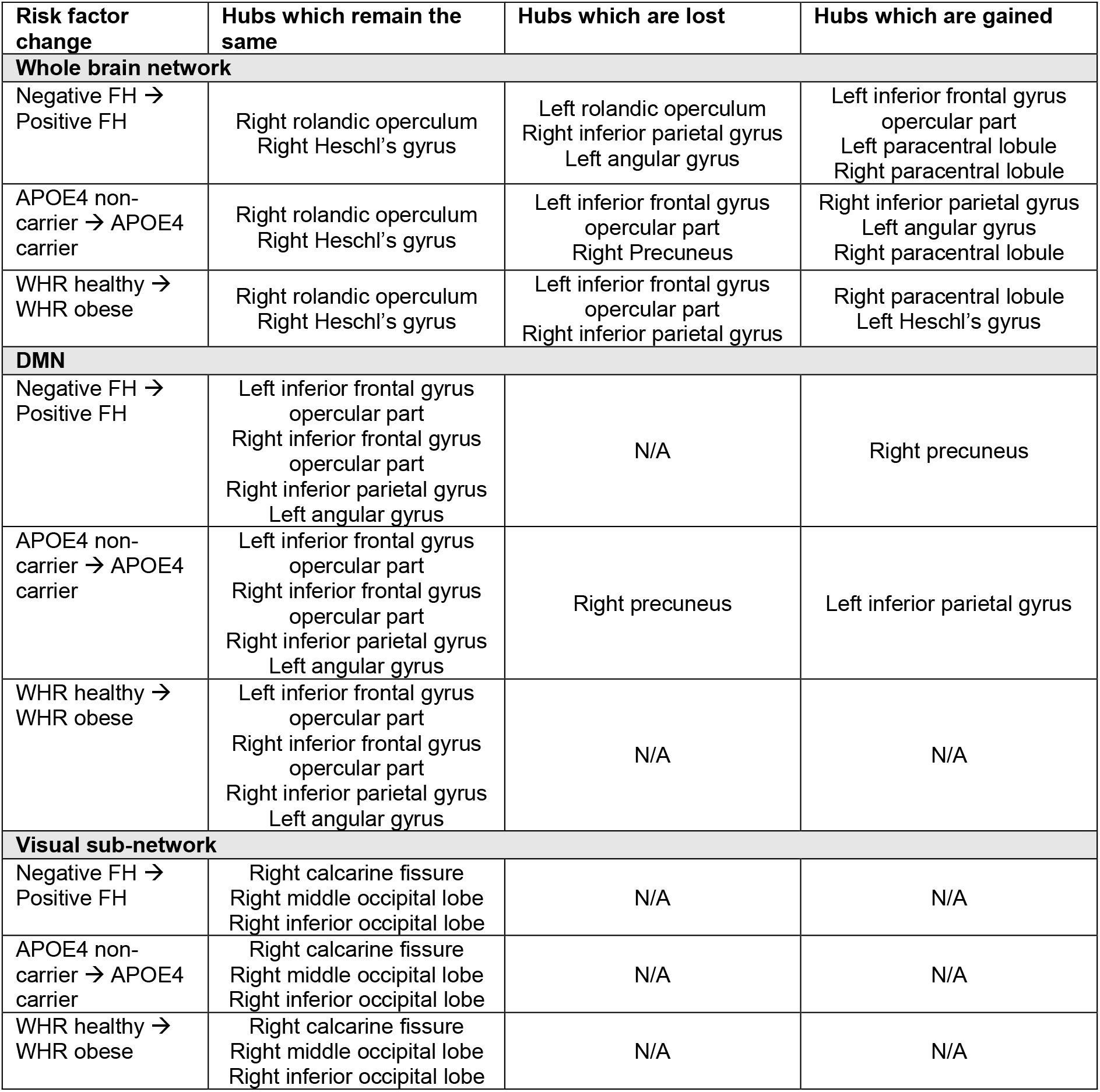
Hub changes as a function of risk factor. *For the whole brain analysis (top)*: The right rolandic operculum and right Heschl’s gyrus remain constant when switching from a low-risk - no family history of dementia (FH), no Apolipoprotein ε4 (APOE4) allele, healthy waist-hip ratio (WHR) score - to a high-risk group (FH, APOE4 or obese). Whereas the right paracentral lobule is consistently gained. Furthermore, a few more hubs are gained or lost, albeit inconsistent across risk factor groups. *In the DMN analysis (middle)*: Individuals with a FH had a hub in the right precuneus in contrast to those without. Conversely this hub is lost between individuals with no APOE4 in comparison to those that carry APOE4 and instead a hub is gained within left inferior parietal gyrus. *In the visual sub-network analysis (bottom)*: Hubs within the Right calcarine fissure Right middle occipital lobe Right inferior occipital lobe remained in all risk factor manipulations.

| Risk factor change | Hubs which remain the same | Hubs which are lost | Hubs which are gained |
| --- | --- | --- | --- |
| **Whole brain network** |  |  |  |
| Negative FH → Positive FH | Right rolandic operculum<br>Right Heschl's gyrus | Left rolandic operculum<br>Right inferior parietal gyrus<br>Left angular gyrus | Left inferior frontal gyrus opercular part<br>Left paracentral lobule<br>Right paracentral lobule |
| APOE4 non-carrier → APOE4 carrier | Right rolandic operculum<br>Right Heschl's gyrus | Left inferior frontal gyrus opercular part<br>Right Precuneus | Right inferior parietal gyrus<br>Left angular gyrus<br>Right paracentral lobule |
| WHR healthy → WHR obese | Right rolandic operculum<br>Right Heschl's gyrus | Left inferior frontal gyrus opercular part<br>Right inferior parietal gyrus | Right paracentral lobule<br>Left Heschl's gyrus |
| **DMN** |  |  |  |
| Negative FH → Positive FH | Left inferior frontal gyrus opercular part<br>Right inferior frontal gyrus opercular part<br>Right inferior parietal gyrus<br>Left angular gyrus | N/A | Right precuneus |
| APOE4 non-carrier → APOE4 carrier | Left inferior frontal gyrus opercular part<br>Right inferior frontal gyrus opercular part<br>Right inferior parietal gyrus<br>Left angular gyrus | Right precuneus | Left inferior parietal gyrus |
| WHR healthy → WHR obese | Left inferior frontal gyrus opercular part<br>Right inferior frontal gyrus opercular part<br>Right inferior parietal gyrus<br>Left angular gyrus | N/A | N/A |
| **Visual sub-network** |  |  |  |
| Negative FH → Positive FH | Right calcarine fissure<br>Right middle occipital lobe<br>Right inferior occipital lobe | N/A | N/A |
| APOE4 non-carrier → APOE4 carrier | Right calcarine fissure<br>Right middle occipital lobe<br>Right inferior occipital lobe | N/A | N/A |
| WHR healthy → WHR obese | Right calcarine fissure<br>Right middle occipital lobe<br>Right inferior occipital lobe | N/A | N/A |

### 2.6 Analysis of internally important nodes/hubs in the DMN

Hubs were identified within the left and right opercular parts of the inferior frontal gyrus, right inferior parietal gyrus and left angular gyrus in individuals with no FH, whereas individuals with FH had hubs within the left and right opercular parts of the inferior frontal gyrus, right inferior parietal gyrus, left angular gyrus and right precuneus. Both individuals of healthy weight and individuals who were obese had hubs within the left and right opercular parts of the inferior frontal gyrus, right inferior parietal gyrus and left angular gyrus. In participants without the APOE4 allele, hubs were identified in the left and right opercular parts of the inferior frontal gyrus, right inferior parietal gyrus, left angular gyrus and right precuneus, whereas individuals who carry APOE4 had hubs within the left and right opercular part of the inferior frontal gyrus, left and right inferior parietal gyrus and left angular gyrus.

As opposed to the analysis of the whole brain, there were no consistent changes of hubs within the DMN as a result of risk factor profile. Individuals without a FH compared to those with a FH gained a hub within the right precuneus whereas this hub was lost in the transition between APOE4 non-carriers to carriers and instead they gained a hub in the left inferior parietal gyrus (Table 8).

### 2.7 Analysis of internally important nodes/hubs in the visual sub-network

Replicator dynamics identified internally important nodes within the right calcarine fissure, right middle occipital lobe and right inferior occipital lobe of the visual sub-network. Each of these hubs were identified regardless of risk factor profile, suggesting AD risk has no effect on hubs within the visual network (Table 8).

## 3. Discussion

To the best of our knowledge, our study investigated for the first time the effects of APOE4 genotypes, central obesity, and family history of dementia on the graph theoretical metrics of structural brain networks derived via tractography, in cognitively healthy adults. The advantage of our analysis methods over conventional structural network analyses lies in the use of integrated structural network matrices which combine, in a data-driven manner, multiple metrics of the white-matter tracts, rather than arbitrarily using one metric. This means that more information is included in the individual structural network matrices.

Graph theoretical metrics expressing segregation and integration of each participant’s structural brain connectome were calculated for the whole-brain and for two sub-networks, the DMN (which is known to be impaired in AD) and the visual network (used here as a control network). Multivariate analyses revealed no significant effects for either whole-brain or for the sub-networks, which suggests that there are no differences in network measures for any of the risk factors (APOE4, family history of dementia or central obesity). This interesting finding, which indicates that the integration and segregation properties of these structural networks are preserved in asymptomatic individuals at heightened risk of developing AD, could point to a possible compensatory mechanism that leads to minimal functional disruption (as indicated by the normal cognitive abilities of our sample). We note, however, that it is not known when, or indeed if, any of these individuals would develop AD. Cortical-thickness based structural brain networks, which reflect different organisational properties to the tractography-derived networks used in our analysis, demonstrated altered properties in subjects with MCI and AD compared to healthy controls following the progress of the disease (Zhou & Liu, 2013). Additionally, Brown et al., 2011 found that tractography-derived structural brain networks in older APOE4 carriers exhibited loss of local interconnectivity in contrast to those of older non-carriers, and that the carriers had impaired memory abilities as well. Finally, Ma et al., 2017 found that structural brain connectivity was disrupted in adults (older than 55 years of age) as a result of an interaction between APOE4 status and developed MCI, more so than it was for APOE4 carriers only. These findings may suggest that structural connectivity changes are not present in cognitively healthy individuals at risk, and reflect a manifestation of established disease and/or of older age.

Looking at the hubs of the whole-brain structural networks of low-risk versus high-risk individuals, we identified that the three subgroups of high-risk individuals (centrally obese, positive FH, and positive APOE4) when compared with individuals in the respective low-risk groups (normal weight, negative FH, and negative APOE4) consistently exhibited a hub in the right paracentral lobule. Importantly, there were no consistent changes of hubs within the DMN and visual network as a result of risk factor profile. The paracentral lobule is located on the medial surface of the cerebral hemisphere and includes parts of both the frontal and parietal lobes. It has gyral projections to the medial frontal gyrus, cingulate sulcus, and precuneus and sulcal projections to the paracentral, cingulate, precentral sulci and the pars marginalis of cingulate sulcus. The paracentral lobule controls motor and sensory innervations of the contralateral lower limb. In a recent study, widespread cortical thinning in left hemisphere regions including pericalcarine cortex, supramarginal gyrus, cuneus cortex, lateral occipital cortex, precuneus cortex, fusiform gyrus, superior frontal gyrus, lateral occipital cortex, entorhinal cortex, inferior parietal cortex, isthmus-cingulate cortex, postcentral gyrus, superior parietal cortex, caudal middle frontal gyrus, insula cortex, precentral gyrus and paracentral lobule was observed in patients with AD compared to normal controls (Yang et al., 2019). Another structural MRI study on nondemented aging subjects revealed a modulation of the cortical thickness covariance between the left parahippocampal gyrus and left medial cortex, supplementary motor area, the left medial superior frontal gyrus, and paracentral lobule driven by the interaction of the rs405509 genotype and age (Chen et al., 2015). In our previous genetic risk for dementia study on the same cohort, we explored the impact of these three factors on white matter microstructure (Mole, Fasano, Evans, Sims, Hamilton, et al., 2020). Individuals with the highest genetic risk (FH+ and *APOE*-ε4) showed a reduced macromolecular proton fraction in the right parahippocampal cingulum associated with obesity. However, we observed effects of cortical thickness only on left thalamus (Mole, Fasano, Evans, Sims, Kidd, et al., 2020). Rs405509 is an AD-related polymorphism located in the APOE promoter region that regulates the transcriptional activity of the APOE gene. Abnormal structural brain connectivity was identified between the angular gyrus, superior parietal gyrus, precuneus, posterior cingulum, putamen, precentral gyrus, postcentral gyrus, and paracentral lobule in elders with subjective cognitive decline compared to healthy controls (Kim et al., 2019). These aberrant structural connections were also associated with cognitive scores.

In addition to MRI, PET imaging has identified reduced metabolism in parietal areas in both *APOE*-ε4 carriers with mild cognitive impairment (Paranjpe et al., 2019) and clinical AD (Mosconi et al., 2004). Furthermore, MEG in young healthy *APOE*-ε4 carriers (Koelewijn et al., 2019) has identified hyperconnectivity in right parietal regions, supporting our findings. If the novel phenotype we have identified can potentially predict the development of symptoms in a longitudinal study of the same cohort, it could be used as an early biomarker of dementia.

### Assessment of our analysis

Our findings would benefit from replication in a larger sample due to the fragmentation of the initial sample into subgroups with the different risk profiles. It would also be beneficial for structural network analyses to include measures which are believed to play a more important role in the functional performance of the brain, such as myelination of the white matter tracts (Messaritaki et al., 2021) and axonal diameter. We finally note that the thresholding of structural connectivity matrices derived from tractography is still issue of debate. Buchanan et al., 2020; Civier et al., 2019; Drakesmith et al., 2015 have shown the possible effects of thresholding when different tractography methods are used. In our analysis, we adopted a modest thresholding of 5 streamlines, to reduce possible false positives.

## 4. Conclusion

In conclusion, our study did not detect any changes in structural brain networks that would imply alterations in the integration and segregation structural network properties in cognitively healthy individuals with different risk factors. We identified the right paracentral lobule as a hub brain area in high-risk individuals but not in low-risk individuals. A longitudinal study of the same cohort with the incorporation of functional neuroimaging data could evaluate this phenotype further.

## Authorship confirmation statement

- HC ran the analysis and drafted the methods and results sections.
- EM provided MATLAB code for the analysis, contributed to study design, and drafted parts of the manuscript.
- SID provided MATLAB code for the analysis, contributed to study design and drafted the manuscript.
- CMB was responsible for the conceptualization of CARDS, draft review & editing, and funding acquisition.

All authors approved the final version of the manuscript.

## Footnotes

1 https://github.com/stdimitr/integrated_structural_brain_networks

2 https://github.com/stdimitr/Network_Metrics

3 https://github.com/stdimitr/consistent_hubs_cohort

