## Supplementary Figures for "Dementia risk factors modify hubs but leave other connectivity measures unchanged in asymptomatic individuals: a graph theoretical analysis"

### S1: Residual distributions before and after data cleaning: whole-brain network

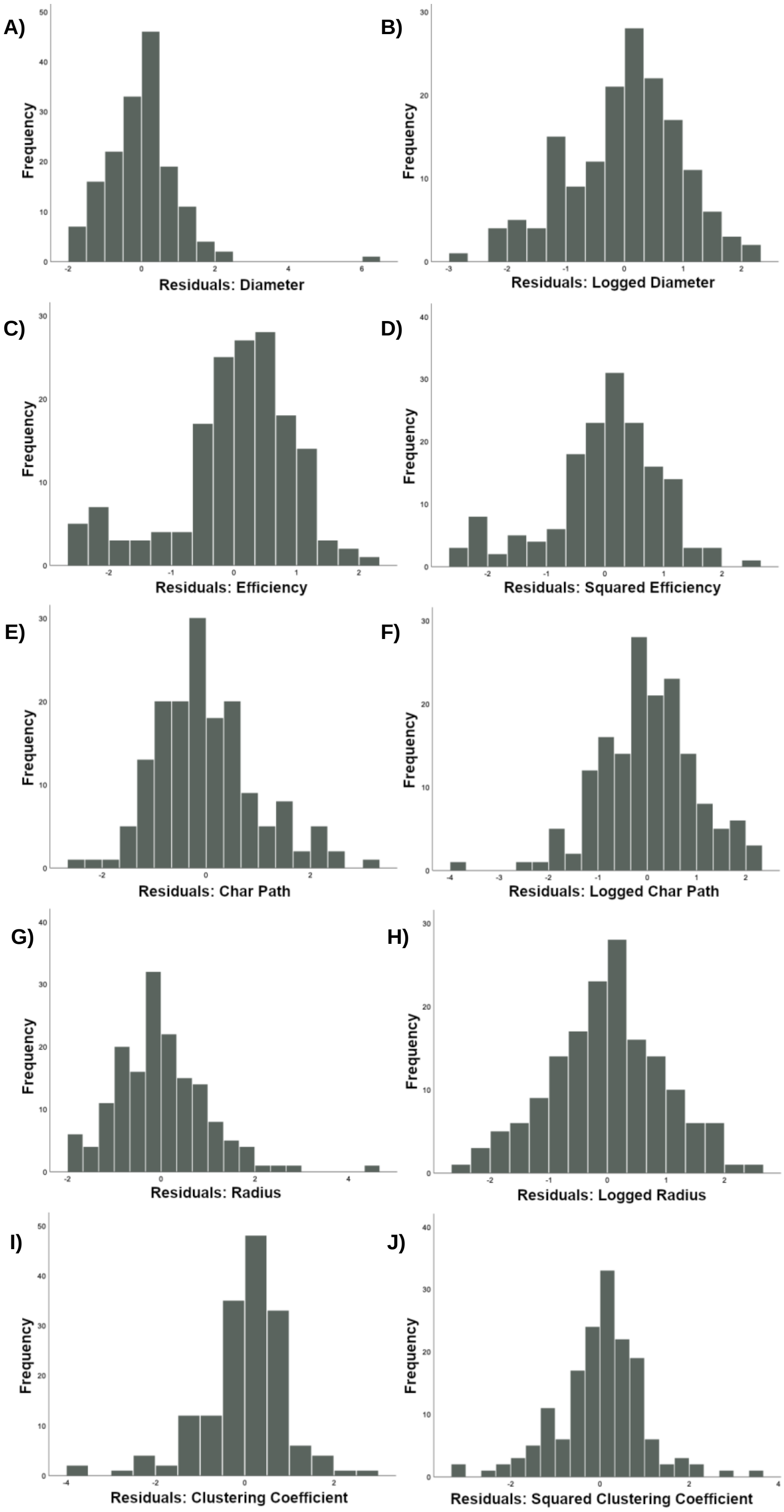

**Figure S1. Histograms of standardized residuals for the whole-brain analysis.** **A**, **C**, **E**, **G** and **I** represent the data prior to outlier removal and transforms whereas **B**, **D**, **F**, **H** and **J** show the distributions of the cleaned metrics which were subsequently inputted into the final general linear model. **A)** Diameter was logged to reduce skew (**B**). Efficiency (**C**) was squared (**D**). **E)** Characteristic path length (Char path) was logged (**F**). In addition, **G)** radius was log transformed (**H**). Whereas, clustering coefficients (**I**) were squared (**J**).

### S2: Residual distributions before and after data cleaning: DMN sub-network

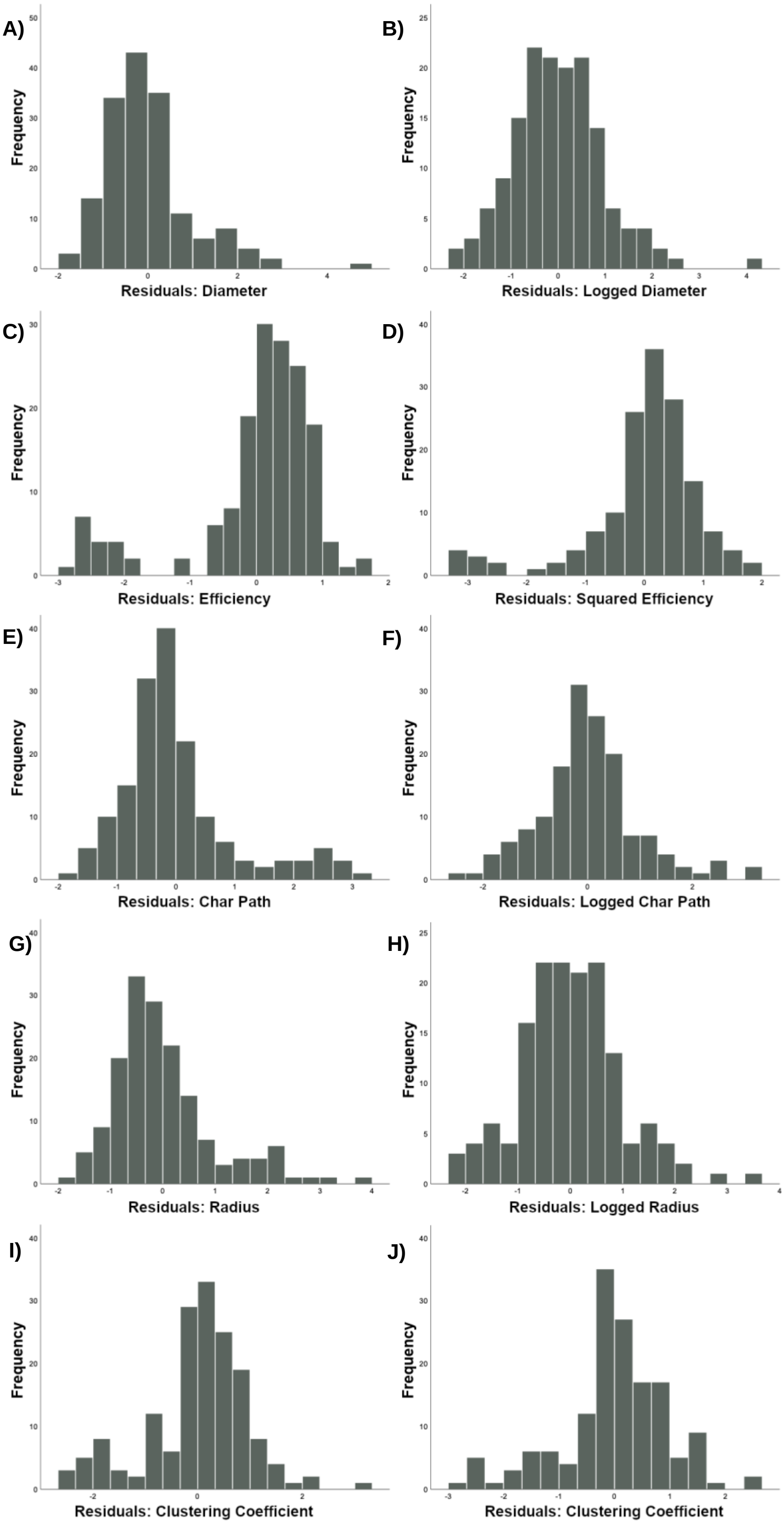

**Figure S2. Histograms of standardized residuals for the analysis of the default mode network.** **A**, **C**, **E**, **G** and **I** represent the raw residuals whereas **B**, **D**, **F**, **H** and **J** show the distributions of the “cleaned” residuals. **A)** Diameter was logged to reduce skew (**B**) and efficiency (**C**) was squared (**D**). **E)** Characteristic path length (Char path) was logged (**F**). In addition, **G)** radius was log transformed (**H**). Whereas, clustering coefficients (**I**) were left un-transformed and only underwent outlier removal (**J**).

### S3: Residual distributions before and after data cleaning: visual sub-network

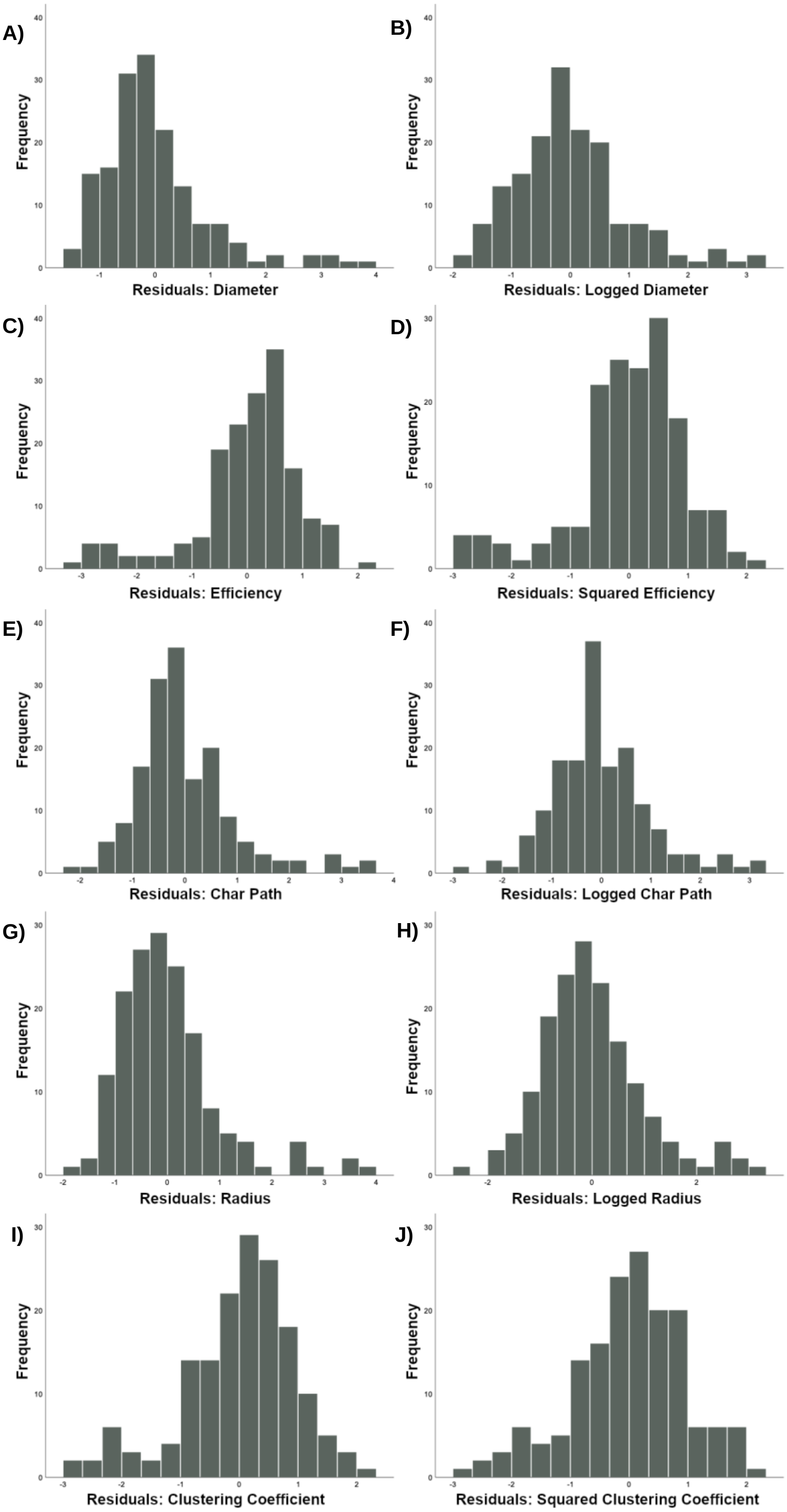

**Figure S3. Distributions of standardized residuals for the visual sub-network analysis.** **A**, **C**, **E**, **G** and **I** illustrate data before outlier removal and transforming to reduce skew whereas **B**, **D**, **F**, **H** and **J** represent the residuals of the “cleaned” data. **A)** Diameter was logged to reduce skew (**B**). Efficiency (**C**) was squared (**D**). **E)** Characteristic path length (Char Path) was logged (**F**). In addition, **G)** radius was log transformed (**H**). Whereas, clustering coefficients (**I**) were squared (**J**) to remove skew.
